## Supplementary Genotypes for "Regeneration following tissue necrosis is mediated by non-apoptotic caspase activity"

**Figure 1.**

(D) hs-FLP/+; hs-P65/ lexAOp-GFP; R85E08-lexADBD, DVE>>GAL4/+

(E) hs-FLP/+; lexAOp-GluR1, hs-P65/lexAOp-GFP; R85E08-lexADBD, DVE>>GAL4/+

(F) +/+; UAS-GFP/+; R85E08GAL4/tubGAL80ts

(G and S1I) +/+; UAS-GFP/UAS-GluR1; R85E08GAL4/tubGAL80ts

(H) +/+; UAS-GFP/+; rn-GAL4/tubGAL80ts

(I and J) +/+; UAS-GFP/UAS-GluR1; rn-GAL4/tubGAL80ts

(K) +/+; UAS-GFP/+; pnr-GAL4/tubGAL80ts

(L and M) +/+; UAS-GFP/UAS-GluR1; pnr-GAL4/tubGAL80ts

(N) +/+; UAS-GFP/+; R73G07-GAL4/tubGAL80ts

(O and P) +/+; UAS-GFP/UAS-GluR1; R73G07-GAL4/tubGAL80ts

(Q and T) +/+; UAS-GFP/+; hh-GAL4/tubGAL80ts

(R, R’, S, S’,T, and S1I) +/+; UAS-GFP/UAS-GluR1; hh-GAL4/tubGAL80ts

**Supplementary figure related to Figure 1.**

(A) +/+; nub-GAL4, UAS-GFP/UAS-GluR1; tubGAL80ts/+

(B) hs-FLP/+; lexAop-GluR1, hs-P65/ +; R85E08-lexADBD, DVE>>GAL4/+

(C and F) +/+; ptc-GAL4, UAS-RFP/+; tubGAL80ts/+

(D, D’, E, E’, F and I) +/+; ptc-GAL4, UAS-RFP/UAS-GluR1; tubGAL80ts/+

(G) hs-FLP; +/+; act>>GAL4, UAS-RFP/+

(H, H’,H’’, I and J) hs-FLP; UAS-GluR1/+; act>>GAL4, UAS-RFP/tubGAL80ts

**Figure 2.**

(A) hs-FLP/+; hs-P56/+ ; R85E08-lexADBD, DVE>>GAL4/10xSTAT-GFP

(B) hs-FLP/+; lexAOp-GluR1, hs-P65/+ ; R85E08-lexADBD, DVE>>GAL4/10xSTAT-GFP

(C and E) hs-FLP/+; lexAOp-GluR1, hs-P65/ UAS-yRNAi; R85E08-lexADBD, DVE>>GAL4/+

(D and E) hs-FLP/+; lexAOp-GluR1, hs-P65/ +; R85E08-lexADBD, DVE>>GAL4/UAS-domeRNAi

(G) hs-FLP/+; lexAOp-GluR1, hs-P65/ UAS-GFP; R85E08-lexADBD, hh-GAL4/ tubGAL80ts

(H and L) hs-FLP/ +; lexAOp-GluR1, hs-P65/ UAS-Stat92ERNAi; R85E080-lexADBD, hh-GAL4/ tubGAL80ts

(I) hs-FLP/ +; hs-P56/ UAS-yRNAi ; R85E08-lexADBD, DVE>>GAL4/+

(J and L) hs-FLP/ +; lexAOp-GluR1, hs-P56/ UAS-yRNAi ; R85E08-lexADBD, DVE>>GAL4/ +

(K and L) hs-FLP/+; lexAOp-GluR1, hs-P65/tubGAL80ts; R85E080-lexADBD, hh-GAL4/UAS-wgRNAi

**Supplementary figure related to Figure 2.**

(A) hs-FLP/ +; hs-P65/ lexAOp-GFP; R85E08-lexADBD, DVE>>GAL4/ UAS-domeRNAi
(B) hs-FLP/ +; hs-P65/ lexAOp-GFP; R85E08-lexADBD, DVE>>GAL4/ UAS-hop48A
(C) hs-FLP/ +; lexAOp-GluR1, hs-P65/ lexAOp-GFP; R85E08-lexADBD, DVE>>GAL4/ UAS-hop48A
(D) hs-FLP/ +; lexAOp-GluR1, hs-P65/ UAS-GFP; R85E080-lexADBD, hh-GAL4/ tubGAL80ts
(E) (H) +/+; +/+; hh-GAL4/10xSTAT-GFP
(F) (I) +/+; UAS-Stat92ERNAi/CyO; hh-GAL4/10xSTAT-GFP

(G) +/+; UAS-GFP/UAS-Stat92ERNAi; hh-GAL4/tubGAL80ts

(H) +/ +; UAS-GFP/ tubGAL80ts; hh-GAL4/ UAS-wgRNAi

(I) +/ +; UAS-GFP/ tubGAL80ts; hh-GAL4/ UAS-Zfh2RNAi

(J and Figure 2L) hs-FLP/+; lexAOp-GluR1, hs-P65/tubGAL80ts; R85E080-lexADBD, hh-GAL4/UAS-Zfh2RNAi

(K) vgQE-lacZ/+

(L) hs-FLP/ +; lexAOp-GluR1, hs-P65/ +; R85E08-lexADBD, DVE>>GAL4/ vgQE-lacZ

(M) +/ +; UAS-Wg/ +; hh-GAL4/ tubGAL80ts

(N) hs-FLP/ +; lexAOp-GluR1, hs-P65/ UAS-Wg; R85E080-lexADBD, hh-GAL4/ tubGAL80ts

**Figure 3.**

(A-A’’’) hs-FLP/ +; lexAOp-GluR1, hs-P65/ +; R85E08-lexADBD, DVE>>GAL4/ PCNA-GFP

(B-B’’’, E, F, G, K and O) hs-FLP/ +; lexAOp-GluR1, hs-P65/ UAS-yRNAi; R85E08-lexADBD, DVE>>GAL4/ +

(C-C’’’) hs-FLP/ +; lexAOp-hepCA, hs-P65/ UAS-yRNAi; R85E08-lexADBD, DVE>>GAL4/ +

(D-D’’’, F, H, L and O) hs-FLP/ +; lexAOp-GluR1, hs-P65/ UAS-mir(RHG); R85E08-lexADBD, DVE>>GAL4/ +

(I, M and O) hs-FLP/ +; lexAOp-GluR1, hs-P65/ UAS-mir(RHG); R85E08-lexADBD, DVE>>GAL4/ DRWNT-GAL80

(J, N and O) w/ +; lexAOp-GluR1, hs-P65/ UAS-mir(RHG); R85E08-lexADBD/ R85E08-GAL4

**Supplementary figure related to Figure 3.**

(A and C) hs-FLP/ +; hs-P65/ +; R85E08-lexADBD, DVE>>GAL4/ PCNA-GFP

(B and C) hs-FLP/ +; lexAOp-GluR1, hs-P65/ +; R85E08-lexADBD, DVE>>GAL4/ PCNA-GFP

(D-D’’’, R and G) hs-FLP/ +; hs-P65/ UAS-yRNAi; R85E08-lexADBD, DVE>>GAL4/ +

(F and G) hs-FLP/ +; hs-P65/ UAS-mir(RHG); R85E08-lexADBD, DVE>>GAL4/ +

**Figure 4.**

(A and A’) hs-FLP/ +; hs-P65/ wg-lacZ; R85E08-lexADBD, DVE>>GAL4/ +

(B and B’) hs-FLP/ +; lexAOp-GluR1, hs-P65/ wg-lacZ; R85E08-lexADBD, DVE>>GAL4/ +

(C and C’) hs-FLP/ +; lexAOp-GluR1, hs-P65/ wg-lacZ; R85E08-lexADBD, DVE>>GAL4/ UAS-P35

(D and D’) hs-FLP/ +; hs-P65/ +; R85E08-lexADBD, DVE>>GAL4/ dpp-lacZ

(E and E’) hs-FLP/ +; lexAOp-GluR1, hs-P65/ + ; R85E08-lexADBD, DVE>>GAL4/ dpp-lacZ

(F and F’) hs-FLP/ +; lexAOp-GluR1, hs-P65/ UAS-P35; R85E08-lexADBD, DVE>>GAL4/ dpp-lacZ

(G) hs-FLP/ +; lexAOp-GluR1, hs-P65/ AP-1-GFP; R85E08-lexADBD, DVE>>GAL4/ +

(H) hs-FLP/ +; lexAOp-GluR1, hs-P65/ UAS-Cat; R85E08-lexADBD, DVE>>GAL4/ UAS-Sod1

(I) hs-FLP/ +; lexAOp-GluR1, hs-P65/+; R85E08-lexADBD, DVE>>GAL4/ UAS-DuoxRNAi

(J and M) hs-FLP/ +; lexAOp-GluR1, hs-P65/ UAS-yRNAi; R85E08-lexADBD, DVE>>GAL4/ +

(K and M) hs-FLP/ +; lexAOp-GluR1, hs-P65/ UAS-P35; R85E08-lexADBD, DVE>>GAL4/ +

(L and M) hs-FLP/ +; lexAOp-GluR1, hs-P65/ UAS-P35; R85E08-lexADBD, DVE>>GAL4/ DRWNT-GAL80

**Supplementary figure related to Figure 4.**

(A) hs-FLP/ +; hs-P65/ +; R85E08-lexADBD, DVE>>GAL4/ spi-lacZ

(B) hs-FLP/ +; lexAOp-GluR1, hs-P65/ UAS-P35; R85E08-lexADBD, DVE>>GAL4/ spi-lacZ

(C) hs-FLP/ +; hs-P65/ +; R85E08-lexADBD, DVE>>GAL4/ mol-lacZ

(D) hs-FLP/ +; lexAOp-GluR1, hs-P65/ +; R85E08-lexADBD, DVE>>GAL4/ mol-lacZ

**Figure 5.**

(A-E, G, G’, L and L’) hs-FLP/ +; lexAOp-GluR1, hs-P65/ +; R85E08-lexADBD, DVE>>GAL4/ UAS-GC3Ai

(H, I, M and M’) hs-FLP/ +; lexAOp-GluR1, hs-P65/ DRWNT-GAL80; R85E08-lexADBD, DVE>>GAL4/ UAS-GC3Ai

(J-K) hs-FLP/ +; lexAOp-hepCA, hs-P65/ +; R85E08-lexADBD, DVE>>GAL4/ UAS-GC3Ai

**Supplementary figure related to Figure 5.**

(A) hs-FLP/ +; hs-P65/ +; R85E08-lexADBD, DVE>>GAL4/ UAS-GC3Ai

(B, E, F, G and H) hs-FLP/ +; lexAOp-GluR1, hs-P65/ +; R85E08-lexADBD, DVE>>GAL4/ UAS-GC3Ai

(C) hs-FLP/ +; lexAOp-GluR1, hs-P65/ DRWNT-GAL80; R85E08-lexADBD, DVE>>GAL4/ UAS-GC3Ai

**Figure 6.**

(A-A’) hs-FLP/ +; lexAOp-GluR1, hs-P65/ +; R85E08-lexADBD, DVE>>GAL4/ DBS-GFP

(B-B’’’) w/ +; lexAOp-GluR1, hs-P65/ CasExpress, tubGAL80ts; R85E08-lexADBD/ UAS-RFP, UAS-FLP, ubi>>stinger (G-TRACE)

(C) w/ +; lexAOp-hepCA, hs-P65/ CasExpress, tubGAL80ts; R85E08-lexADBD/ UAS-RFP, UAS-FLP, ubi>>stinger (G-TRACE)

(D and E) w/ +; lexAOp-GluR1, hs-P65/ CasExpress; R85E08-lexADBD/ UAS-GFP

**Supplementary figure related to Figure 6.**

(A) w/ +; lexAOp-GluR1, hs-P65/ CasExpress, tubGAL80ts; R85E08-lexADBD/ UAS-RFP, UAS-FLP, ubi>>stinger (G-TRACE)

**Figure 7.**

(A-B, K and L) hs-FLP/ +; lexAOp-GluR1, hs-P65/ UAS-yRNAi; R85E08-lexADBD, DVE>>GAL4/ +

(C-D, K and L) hs-FLP/ +; lexAOp-GluR1, hs-P65/ UAS-mir(RHG); R85E08-lexADBD, DVE>>GAL4/ +

(E-F, K, L and M) hs-FLP/ +; lexAOp-GluR1, hs-P65/ +; R85E08-lexADBD, DVE>>GAL4/ droncI29

(G-H, K and L) hs-FLP/ +; lexAOp-GluR1, hs-P65/ UAS-dronc-CARD; R85E08-lexADBD, DVE>>GAL4/ +

(I-J, K and L) hs-FLP/ +; lexAOp-GluR1, hs-P65/ +; R85E08-lexADBD, DVE>>GAL4/ UAS-DIAP1

(M) hs-FLP/ w1118; lexAOp-GluR1, hs-P65/ +; R85E08-lexADBD, DVE>>GAL4/ +

**Supplementary figure related to Figure 7.**

(A) hs-FLP/ w1118; hs-P65/ +; R85E08-lexADBD, DVE>>GAL4/ +

hs-FLP/ +; hs-P65/ +; R85E08-lexADBD, DVE>>GAL4/ droncI29

hs-FLP/ w1118; lexAOp-GluR1, hs-P65/ +; R85E08-lexADBD, DVE>>GAL4/ +

hs-FLP/ +; lexAOp-GluR1, hs-P65/ +; R85E08-lexADBD, DVE>>GAL4/ droncI29
